# IL-13-programmed airway tuft cells produce PGE2, which promotes CFTR-dependent mucociliary function

**DOI:** 10.1101/2022.05.11.491556

**Authors:** Maya E. Kotas, Camille M. Moore, Jose G. Gurrola, Steven D. Pletcher, Andrew N. Goldberg, Raquel Alvarez, Sheyla Yamato, Preston E. Bratcher, Ciaran A. Shaughnessy, Pamela L. Zeitlin, Irene Zhang, Yingchun Li, Michael T. Montgomery, Keehoon Lee, Emily K. Cope, Richard M. Locksley, Max A. Seibold, Erin D. Gordon

**Author notes:** Corresponding authors: Erin D. Gordon, MD, Division of Pulmonary, Critical Care, Allergy and Sleep Medicine, Department of Medicine, University of California, San Francisco, 513 Parnassus Ave, HSE 201 San Francisco, CA 94143, Max A. Seibold, PhD, Center for Genes, Environment, and Health National Jewish Health, University of Colorado-AMC, School of Medicine, Smith Building; A648, Denver, CO 80206, National Jewish Health, 1400 Jackson Street, A642, Denver, CO 80206. equal contribution (Maya E. Kotas, Camille M. Moore).

## Abstract

Chronic type 2 (T2) inflammatory diseases of the respiratory tract are characterized by mucus overproduction and disordered mucociliary function, which are largely attributed to the effects of IL-13 on common epithelial cell types (mucus secretory and ciliated cells). The role of rare cells in airway T2 inflammation is less clear, though tuft cells have been shown to be critical in the initiation of T2 immunity in the intestine. Using bulk and single cell RNA sequencing of airway epithelium and mouse modeling, we find that IL-13 expands and programs airway tuft cells towards eicosanoid metabolism, and that tuft cell deficiency leads to a reduction in airway prostaglandin E2 (PGE2) concentration. Allergic airway epithelia bear a signature of prostaglandin E2 activation, and PGE2 activation leads to CFTR-dependent ion and fluid secretion and accelerated mucociliary transport. Together these data reveal a role for tuft cells in regulating epithelial mucociliary function in the allergic airway.

## Introduction

Chronic rhinosinusitis with nasal polyps (CRSwNP) and asthma are common airway diseases characterized by persistent type 2 (T2), or “allergic” inflammation. The two diseases share a number of genetic risk loci (1, 2) and patients with disease affecting the upper airway have more severe lower airway symptoms, consistent with overlapping pathological mechanisms (3). A key finding in both conditions is the IL-13-induced shift in epithelial cell type composition in favor of mucus secretory cells (goblet cells) with resultant pathologic secretions. While IL-13 has also been shown to alter the transcriptional profile and function of all common airway epithelial cell types (basal, secretory, ciliated) (4), functional shifts in rare cells are incompletely described.

The airway epithelium is composed of prevalent cell types such as basal, ciliated, and secretory/mucus cells, interspersed with rare cell types such as neuroendocrine cells, ionocytes, and tuft cells. These rare cells, while technically challenging to interrogate, are increasingly recognized for their contributions to airway homeostasis and mucosal immunity. For example, the discovery of ionocytes in human and mouse trachea revealed that these are the highest expressers of cystic fibrosis transmembrane receptor (CFTR), an ion channel important in regulating the composition of airway surface liquid and maintaining bacterial host defense (5, 6). Neuroendocrine cells can sense and protect against hypoxia (7), as well as augment type 2 inflammation through effects on ILC2s (8). Tuft cells initiate and amplify type 2 inflammation in the mouse intestine (9–11), and can stimulate protective reflexes such as cough or apnea, as well as inflammation, in the airway (12, 13).

The full range of tuft cell outputs and functions, particularly in human disease, remains poorly understood. In addition to IL-25, mouse tuft cells are capable of producing leukotrienes, prostaglandins, and acetylcholine, suggesting potentially pleiotropic roles in tissue homeostasis. Though recent studies have suggested that tuft cells may be a source of IL-25 in nasal polyposis (14, 15) and activate type 2 immunity as in the mouse intestine, perturbations of tuft cells in human airway disease have not been fully explored. Existing atlases of the cellular landscape in CRSwNP and allergic asthma that have broadly examined immune, stromal, and epithelial cells have not identified these rare but potentially important cells (16, 17). With this in mind, we sought to identify and phenotype tuft cells in the human respiratory epithelium, and to explore their role in the pathobiology of allergic airway disease.

## Results

### Tuft cells expand and adopt an “allergic” transcriptional program of eicosanoid metabolism in nasal polyps

To define the cellular composition of human allergic (T2 inflamed) sinus tissue, we collected epithelial brushes from ethmoid-based polyps of 5 patients with CRSwNP or from healthy ethmoid tissue of 4 controls (Supplemental Table 1) and subjected dissociated cells to single-cell RNA sequencing (scRNA-Seq). Initial clustering of 116,358 epithelial cells (Supplemental Figure 1A) revealed 15 distinct clusters, each including cells from all patients (Supplemental Table 2). Cluster identities were assigned based on expression of previously described cell type markers (Supplemental Figure 1B) and then aggregated based on similar expression profiles into 10 cell types: basal, proliferating basal, “hillock” basal (6, 17), supra-basal, early secretory, secretory, mucus secretory (goblet), ciliated, and a rare cell type cluster containing markers of both ionocytes and tuft cells (Figure 1A, Supplemental Table 3). We found a similar percentage of most cell types between polyp and control patients suggesting a largely preserved cellular composition (Supplemental Figure 1C). We did, however, observe an overrepresentation of mucus secretory (goblet) cells with high expression of both *MUC5AC* and *MUC5B* in polyp epithelium, consistent with the mucus metaplasia observed in patients with CRSwNP (18) (Supplemental Figure 1C, Supplemental Table 2).

**Figure 1.**
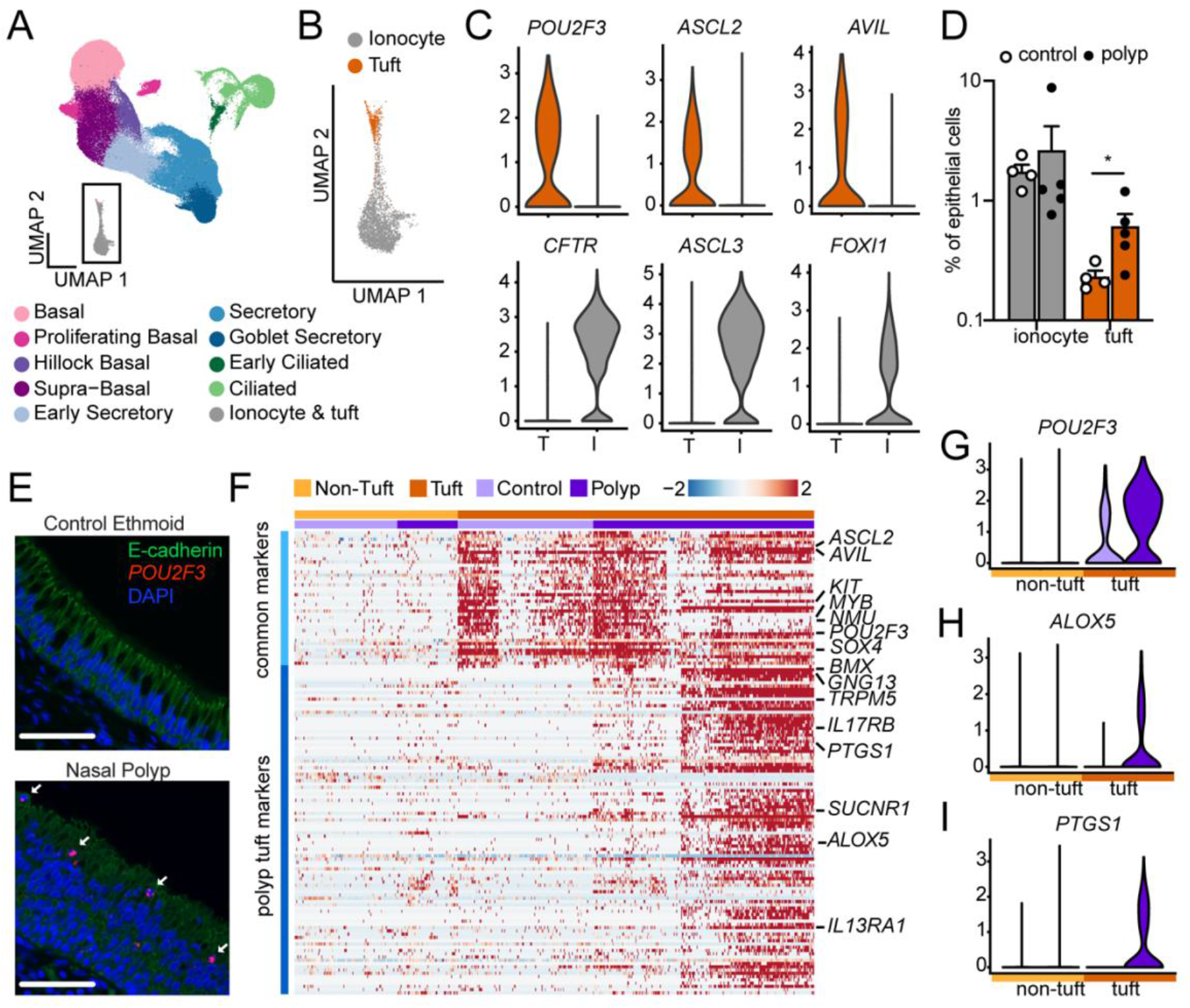
Single cell sequencing reveals expansion and allergic activation of tuft cells in nasal polyps. **(A)** Uniform manifold approximation and projection (UMAP) of scRNA-seq of epithelial cells from healthy ethmoid sinus (control; n=4) or nasal polyp (n=5) (total cells = 116,358) reveals 10 cell types. **(B)** Expanded view of inset in (A) showing tuft cells and ionocytes identified by hierarchical sub-clustering. **(C)** Expression of tuft cell and ionocyte marker genes in tuft and ionocyte sub-clusters. **(D)** Percentage of ionocytes and tuft cells among total epithelial cells in control or polyp. error bars indicate mean +/- SEM. *p<0.05 by Mann-Whitney t test with correction for multiple comparisons. **(E)**RNA *in situ* hybridization for *POU2F3* (red) and immunofluorescence for E-cadherin (green) identifies increased numbers of tuft cells (arrow) in nasal polyp epithelium as compared to control ethmoid (representative of 3 samples from 3 patients of each type). **(F)** Shared tuft cell marker genes (“common markers”) and DEGs (“polyp tuft markers”) in tuft (dark orange) and non-tuft cells (light orange) from control (light purple) (dark purple) epithelium. **(G)** Expression of representative common tuft cell marker *POU2F3* and polyp tuft markers **(H)** *ALOX5* and **(I)** *PTGS1*.

Subclustering of the rare cell type cluster identified both tuft cells (expressing *POU2F3*, *ASCL2* and *AVIL*) and ionocytes (expressing *CFTR*, *ASCL3*, and *FOXI1*) (Figure 1B-C, Supplemental Figure 1D), whereas neuroendocrine cells were not found. The cytokine *IL25*, characteristically produced by mouse tuft cells, was not detectable in human tuft cells, consistent with prior reports (19, 20). While ionocyte frequencies did not differ with disease (median 1.7 vs 1.2%, Figure 1D, Supplemental Table 2), we observed approximately 2.5-fold more tuft cells in polyp compared to healthy sinus (0.2% vs 0.53%, Figure 1D, Supplemental Table 2). The presence of tuft cells and ionocytes (Supplemental Figure 1E) and the increase in tuft cells (Figure 1E) were confirmed by *in situ* RNA hybridization in histological sections from control and nasal polyp tissue.

We next explored whether the expression profile of tuft cells was altered in polyp patients. Comparing tuft cells to all other cell types, we identified a set of tuft cell markers common to both polyp patients and controls, including *ASCL2*, *POU2F3*, and *AVIL*, which we call “common markers.” Then, comparing tuft cells from control vs. polyp patients, we identified a second set of genes increased in the tuft cells of polyp patients, including *BMX*, *GNG13*, *TRPM5*, and *PTGS1*, which we call “polyp tuft markers” (Figure 1F). All of the tuft cells from healthy subjects and a subset of those from polyp patients expressed only common tuft markers. The remaining tuft cells in polyp patients, which we called “allergic” tuft cells, expressed both common and polyp tuft markers. Polyp tuft markers were uniquely enriched for genes associated with the arachidonic acid metabolic pathway including both *PTGS1* and *ALOX5* (Fig. 1G-I, Supplemental Figure 1F-G, Supplemental Figure 2A-B. We compared our two tuft cell phenotypes (healthy and allergic) to publicly available tuft cell transcriptomic profiles. Healthy human sinus tuft cells resembled previous descriptions of human airway tuft cells (19, 20), while canonical markers of mouse tuft cells (21) such as *TRPM5* and *DCLK1* were present only in the allergic tuft cell population in polyp patients (Supplemental Figure 2C). Markers for previously described subsets of small intestinal or airway mouse tuft cells (6, 22) did not discriminate between healthy and allergic human airway tuft cells (Supplemental Figure 2D).

### Pan-epithelial gene signatures in nasal polyps are imparted by IL-13 and PGE2

CRSwNP is mediated by type 2 inflammation, with a prominent role for IL-13 (16). While many genes have altered expression in the setting of type 2 inflammation, prior work identified a simplified three gene signature of type 2 inflammation in bronchial and nasal epithelium of asthmatics that correlates with clinical phenotype (23, 24). We found that this type 2 gene score was broadly increased across epithelial cell clusters in patients with polyps compared to controls (Supplemental Figure 3A). To build upon this finding, we sought to define the entire repertoire of pan-epithelial gene expression alterations in nasal polyps. We performed differential expression analysis of each cell type, comparing polyp to control groups. We then categorized differentially expressed genes (DEGs) based on the number of epithelial cell types in which they were found (Figure 2A, Supplemental Figure 3B). Most DEGs were only identified in 1-2 epithelial clusters suggesting cell-type-specific responses, but a small subset of DEGs were found in 9 or more epithelial cell types. We defined these DEGs as “pan-epithelial”. Among these DEGs, we identified two co-expressed gene sets (Supplemental Figure 3C). We determined that one set (including *CST1*, *POSTN*, *NTRK2*, and *ALOX15)* was upregulated by IL-13 (Figure 2B), consistent with prior work (4, 23), while the second set of genes (including *PTHLH*, *SLC6A8*, *IGFBP3*, *NDRG1*, *EGLN3*, and *ERO1A*) was not IL-13-inducible (Figure 2B). We reasoned that tuft cell-derived eicosanoids might act on the epithelium to induce this gene signature based on prior reports of arachidonic acid pathway dysregulation in nasal polyposis (25) and our observation of allergic tuft cell expansion. We stimulated human airway epithelial cells with leukotrienes and prostaglandins and found that prostaglandin E2 (PGE2) increased expression of the second gene set (Figure 2C-D, Supplemental Table 7), but not the IL-13-responsive genes.

**Figure 2.**
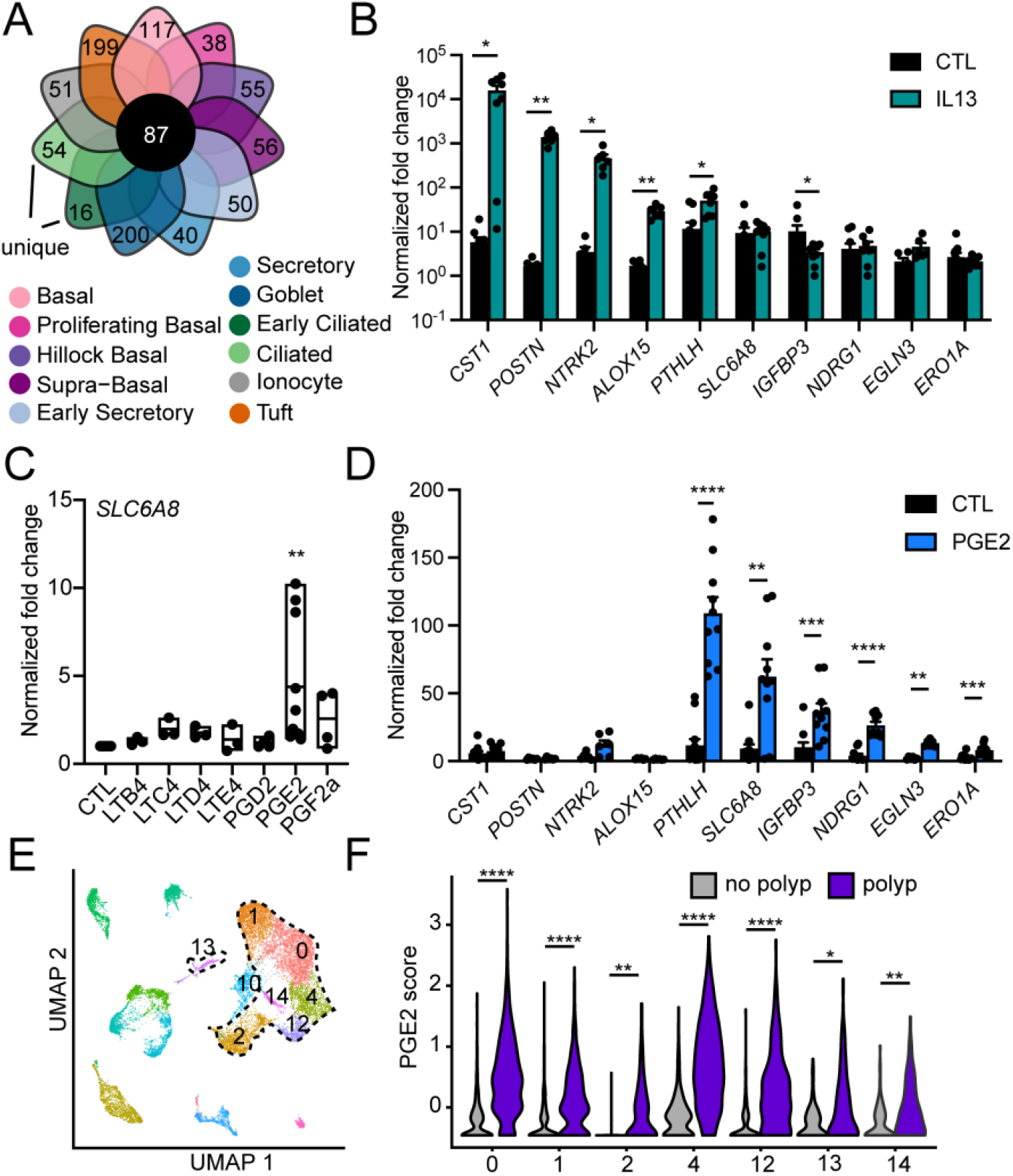
Pan-epithelial gene signatures in nasal polyps are imparted by IL-13 and PGE2. **(A)** Among all differentially expressed genes for each cell type, 87 genes were upregulated in ≥ 9 cell types in polyp epithelium compared to controls and defined as pan-epithelial. **(B)** Fold change in normalized gene expression for tracheal epithelial cells cultured at (ALI) and stimulated with IL-13 (n=10 wells from 6 donors, *p<0.05; **p<0.01; ***p<0.001; ****p<0.0001 by ANOVA with Tukey correction). **(C)**Fold change in normalized *SLC6A8* gene expression in human tracheal epithelial cells cultured at ALI and stimulated with indicated eicosanoids (n=9 donors, **p<0.01 by ANOVA with Dunnett’s correction). Represents similar responses for all genes as shown in (D) **(D)** Fold change in normalized gene expression for tracheal epithelial cells cultured at (ALI) and stimulated with PGE2 (n=10 wells from 6 donors, *p<0.05; **p<0.01; ***p<0.001; ****p<0.0001 by ANOVA with Tukey correction). **(E)** UMAP of scRNA-seq data from surgical sinus tissue of subjects with CRSwNP (“polyp”) or CRSsNP (“no polyp”) whole polyp or non-polyp sinus tissue (16). Epithelial clusters encircled with dashed line. **(F)** PGE2 response gene score in polyp and non-polyp epithelial clusters. *p<0.05; **p<0.01; ***p<0.001; ****p<0.0001 by linear mixed model. Statistical calculations relating to this figure are included in Supplemental Table 7.

To validate our finding of a novel PGE2 gene expression signature in the nasal polyp epithelium, we analyzed published data from single-cell sequencing of whole polyp tissue (16). We re-identified 7 epithelial cell clusters using published markers (Figure 2E) and performed differential expression analysis for each epithelial cell type between samples derived from subjects with or without polyps. The PGE2 activation score was robustly upregulated in all of the epithelial cell type clusters in this independent single-cell dataset (Figure 2F, Supplemental Table 7).

### IL-13 expands and programs airway tuft cells towards PGE2 production

IL-13 is critical to goblet cell metaplasia in type 2 inflammation of the airway, as well as to both tuft and goblet cell expansion in the intestine. To explore the role of IL-13 in tuft cell expansion and PGE2 production in the airway, we induced systemic IL-13 overexpression in mice using hydrodynamic plasmid injection (Figure 3A). Injected mice had high plasma IL-13 levels, whereas circulating IL-13 was not detected in control mice (Supplemental Figure 4A). Tuft cells were increased in both the nasal (Figure 3B-C) and tracheal epithelium (Supplemental Figure 4B-C) following IL-13 overexpression. Single-cell sequencing revealed the emergence of two distinct tuft cell clusters under IL-13 stimulation (Figure 3D-E, Supplemental Figure 4D-E, Supplemental Table 6). Gene expression differences between IL-13-emergent mouse nasal tuft cells versus control mouse tuft cells did not correlate with previously-identified subsets of tuft cells in untreated mouse trachea or intestine (Supplemental Figure 4F). However, the emergent mouse nasal tuft cell clusters expressed genes similar to those observed in the allergic tuft cells of CRSwNP patients, as evident by an increase in the mean expression of the polyp tuft marker genes (i.e. “polyp tuft score”) compared to the tuft cells from unstimulated mice (Figure 3F). Because this mouse model recapitulated key aspects of the tuft cell changes observed in human CRSwNP, we then applied this model to determine if PGE2 production in the respiratory tract was dependent on allergic tuft cells. Using tuft cell-deficient *Pou2f3^-/^*^-^ mice exposed to systemic IL-13, we found a marked reduction in PGE2 metabolites (PGEM) in tracheal lysates of *Pou2f3^-/^*^-^ compared to wildtype mice (Figure 3G-H). PGE2 was also markedly reduced in tracheal organoids generated from *Pou2f3^-/^*^-^ mice, supporting an epithelial cell of origin (Figure 3G, 3I).

**Figure 3.**
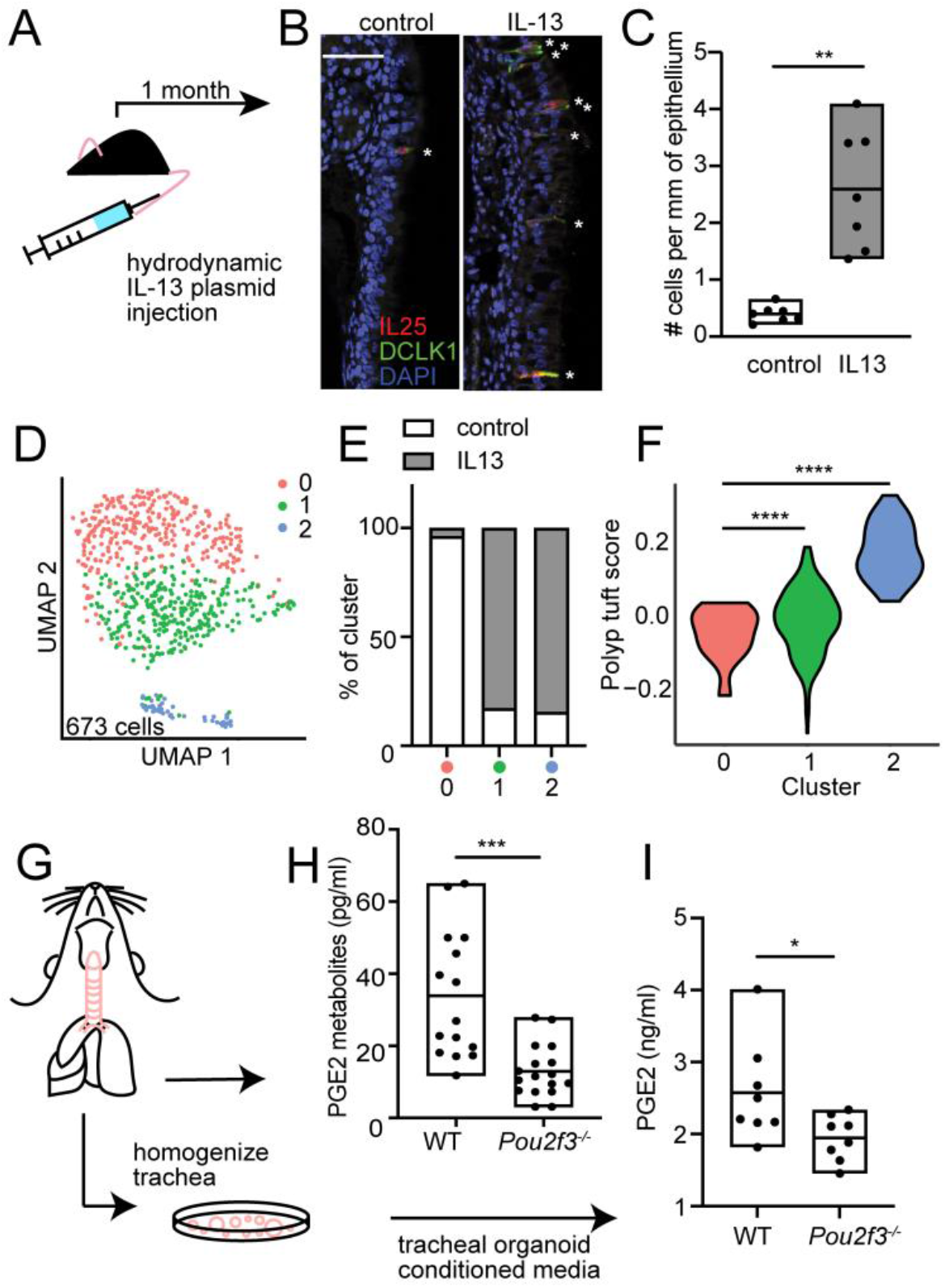
IL-13 expands and programs airway tuft cells towards PGE2 production. **(A)** Type 2 inflammatory mouse model system. **(B)** Representative sections of mouse nasal epithelium after 1 month of IL-13 overexpression or IgG control. Stars mark dual staining IL-25^+^ and DCLK1^+^ tuft cells. Bar indicates 50 µm **(C)** Quantification of tuft cells in nasal epithelium in control and systemic IL-13 expression. **p<0.01 by t-test. **(D)** Subclustering of tuft cells from control or IL-13 overexpressing mouse nasal epithelium. **(E)** Percentage of tuft cells derived from control or systemic IL-13 conditions in each subcluster in (D). **(F)** Human polyp allergic tuft cell gene score in mouse nasal epithelial tuft cell subclusters. ****p<0.0001 by linear regression model. **(G)** Schematic of protocol to measure **(H)** prostaglandin E2 metabolites (PGEM) in whole tracheal tissue from WT or *Pou2f3*^-/-^ mice exposed to systemic IL-13 or **(I)** PGE2 in media from tracheal organoids derived from WT or *Pou2f3*^-/-^ mice. For (H,I), *p<0.05; ***p<0.001 by t-test.

### IL-13-dependent tuft cell programming and PGE2 activation are common features of upper and lower allergic airway disease

To expand our findings across allergic diseases of the upper and lower respiratory tract, we collected epithelial brushings from the sinus of patients with both CRSwNP and asthma (“Polyp”), CRS without asthma or nasal polyps (“CRSsNP”), and control subjects (Supplemental Table 1). Consistent with our single cell sequencing data, epithelial type 2 activation was increased in polyp brushes (Figure 4A), while it was not increased in CRSsNP. In further agreement with our scRNA-Seq data, the “polyp tuft score” was also increased in polyp but not CRSsNP subjects (Figure 4B), and paralleled by an increased in the PGE2 score (Figure 4C-D). Consistent with our finding that IL-13-treated mice deficient in tuft cells have reduced PGE2 in the respiratory tract, and further supporting a causal link between IL-13, allergic tuft cell generation, and PGE2 production, we found a strong correlation between the polyp tuft score and PGE2 activation score, as well as between the type 2 inflammation and epithelial PGE2 activation scores in these subjects (Figure 4D-E).

**Figure 4.**
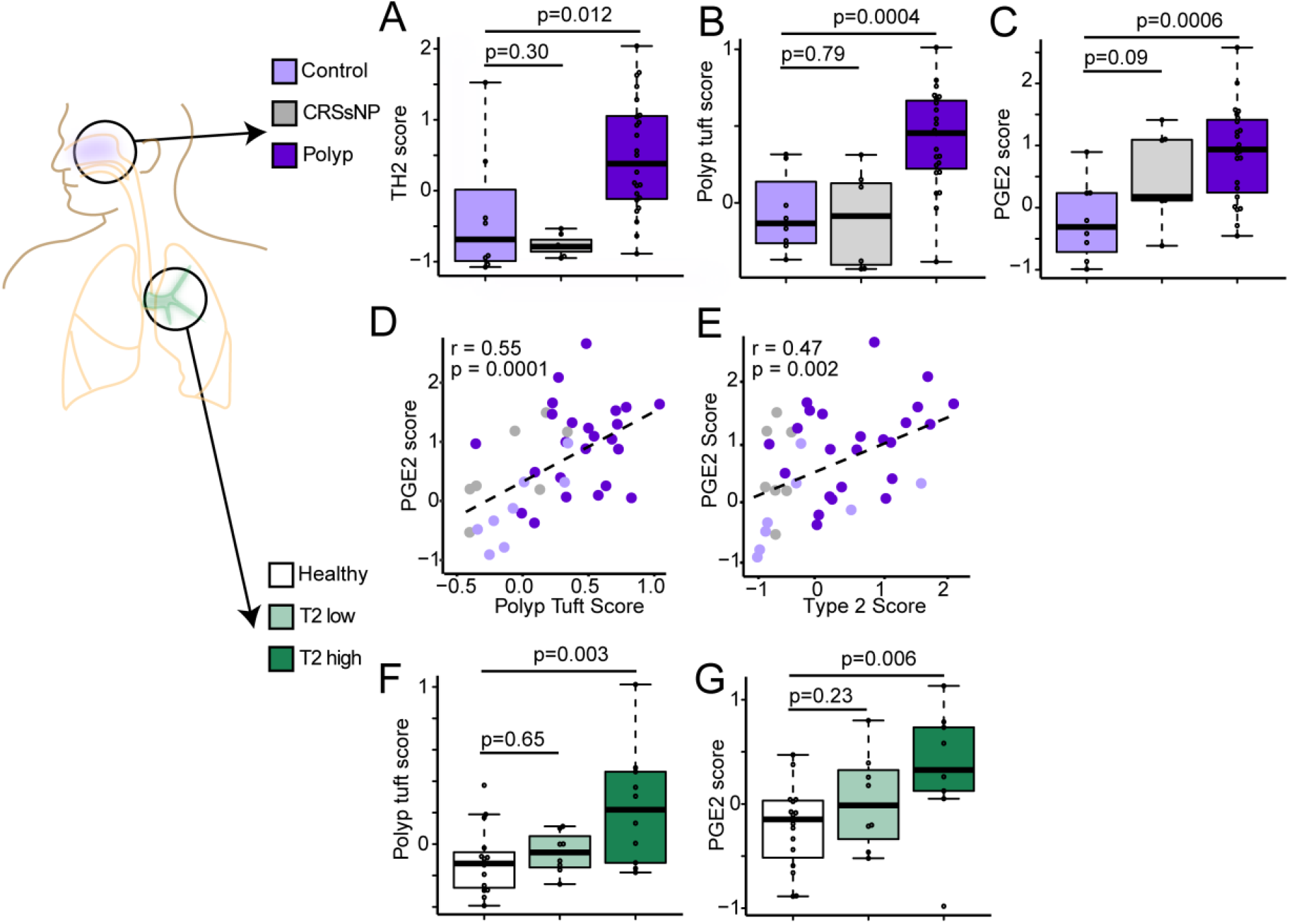
Tuft cell and PGE2 activation are common features of upper and lower allergic airway disease. **(A)** type 2 (3 gene) score, **(B)** polyp tuft score, and **(C)**PGE2 score in RNA sequenced bulk epithelial brushings from sinus tissue of control patients (light purple) or patients with CRS without asthma or nasal polyps (“CRSsNP”; grey), or CRS with asthma and nasal polyps (“Polyp,” dark purple). **(D)** Correlation between PGE2 score and polyp tuft cell score in sinus. **(E)** Correlation between PGE2 score and type 2 score in sinus. For (A-E), n=8 control, n=7 CRSsNP, n=24 polyp subjects, as in Supplementary Table 1. **(F)** Polyp tuft score and **(G)** PGE2 score in RNA sequencing of bulk epithelial brushings the bronchus of healthy patients or asthmatic patients classified as either type 2 low (T2 low) or type 2 high (T2 high). For (F-G), n=16 healthy, n=8 T2 low, n=11 T2 high.

Since chronic IL-13 stimulation in mice led to tuft cell expansion and PGE2 production in both the upper and lower airway, we reasoned that type 2 inflammation may similarly drive tuft cell activation throughout the airway in humans. Indeed, allergic tuft cell transcripts and PGE2 activation were increased in the bronchial epithelium of asthmatics with type 2 inflammation (Figure 4F-G).

### PGE2 regulates epithelial CFTR-dependent fluid secretion and mucociliary transport

To examine the effects of PGE2 on respiratory epithelial function, we cultured healthy human tracheal and sinus epithelium in the presence or absence of PGE2. Of the four PGE2 receptors (EP1-4), EP4 and to a lesser extent EP2 are expressed in airway epithelial cells (26). We found that chronic exposure to PGE2 progressively increased 3D organoid diameter in an EP4- but not EP2-dependent fashion (Figure 5A-B), similar to effects reported in intestinal epithelium (27, 28). Increased organoid diameter in response to PGE2 also occurred in sinus epithelium from polyp patients (Supplemental Figure 5A). Based on their diameter, we estimated organoid surface area increased 1.8-fold with PGE2 treatment, whereas cellular DNA content increased 1.4-fold (Supplemental Figure 5B), reflecting a modest augmentation in cell number and suggesting additional mechanisms of organoid expansion. Moreover, we observed that acute PGE2 application caused organoid swelling over minutes to hours, which was blocked by inhibition of the cystic fibrosis transmembrane receptor (CFTR) (Figure 5C), suggesting organoid swelling was caused by ion and fluid movement. Indeed, PGE2 activated CFTR-dependent currents in 2D cultured human upper airway epithelial cells (Figure 5D). Effective mucociliary transport (MCT) is highly dependent on epithelial ion and fluid secretion, as exemplified by the pathology observed in the disease cystic fibrosis. Consistent with this, we found that PGE2 stimulation increased MCT on the surface of cultured human airway epithelial cells in a CFTR-dependent fashion (Figure 5E).

**Figure 5.**
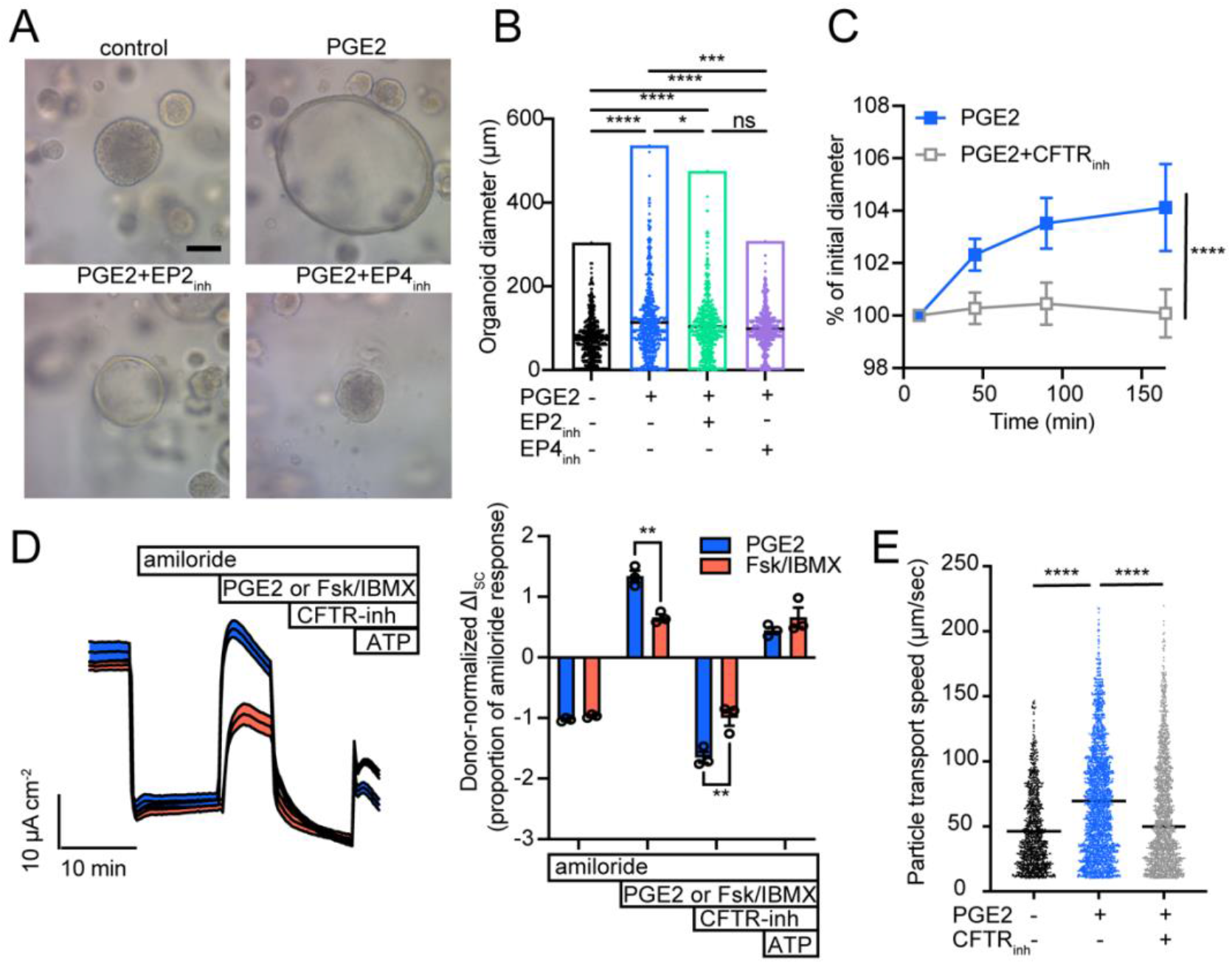
PGE2 regulates epithelial CFTR-dependent fluid secretion and mucociliary transport. **(A)** Human tracheal epithelial organoids cultured for 21 days in the absence or presence of daily stimulation with 1 µg/mL PGE2, or with PGE2 + EP2 inhibitor (EP2_inh_) or PGE2 + EP4 inhibitor (EP4_inh_). 20x magnification; scale bar, 100 µm. representative of at least 2-3 wells per experiment in at least 4 independent experiments with different tracheal and sinus epithelial donors. **(B)** Diameter of organoids in (A). each dot represents 1 of 200 randomly selected organoids per well x 3 replicate wells per condition. Line indicates median diameter. ***p<0.001, *p<0.05, ns = p ≥ 0.05 by 1-way ANOVA with Sidak correction. **(C)** Diameter of chronically PGE2-treated organoids as in (A, B) immediately after acute stimulation with PGE2 +/- CFTR inhibitor 172 (CFTR_inh_). Value +/- 95% CI at each timepoint represents 20 serially-imaged organoids per condition. Representative of 2 independent experiments. ****p<0.0001 by 2-way ANOVA. **(D)** Short circuit current measured in human nasal epithelial cells at ALI after inhibition of epithelial sodium channel-dependent current with amiloride, followed by treatment with PGE2 or cAMP activation by forskolin/IBMX, then by CFTR inhibition, and finally by ATP-dependent activation. Donor normalized quantification of short circuit currents shown in right panel, with each dot representing treatment of an individual epithelial donor. ****p<0.0001 by 2-way ANOVA. **(E)** Particle transport speed on the surface of tracheal epithelial cells cultured at ALI and stimulated with PGE2 +/- CFTR inhibitor 172. Each dot represents one tracked particle on the surface of at least 3 replicate wells from each of 3 donors per condition. ****p<0.0001 by 2-way ANOVA of log-transformed speeds.

## Discussion

Though first observed throughout the conducting airways more than 50 years ago, the rarity of airway tuft cells has been a challenge to their study. As a result, the phenotype and function of these cells has been elusive, and conceptual models have relied heavily on studies in the mouse small intestine. Here we show that tuft cells in the context of allergic diseases of the respiratory tract expand in number and adopt a transcriptional program that augments PGE2 production throughout the airways. PGE2 signaling, in turn, promotes epithelial fluid secretion and mucociliary transport via activation of the CFTR channel. Together, these findings suggest that tuft cells direct mucociliary homeostasis in the allergic airway, and provide insight into their critical homeostatic function.

The foremost purpose of the conducting airways is to defend against particles, microbes, and chemicals while transmitting air to the lung for gas exchange. Mucociliary clearance is the primary airway defense, and therefore critical to its proper function. An abundant body of literature has shown that ionic and fluid movement across the epithelium is critically important in maintaining the function of the airway mucous barrier, impacting the biophysical properties of gel-forming mucins, optimizing the activity of antimicrobial peptides, and enabling mucociliary transit (29–31). These nonredundant roles are exemplified by the consequences of genetic CFTR dysfunction and cystic fibrosis. In the allergic airway, the composition and functional properties of the mucus layer are substantially altered by IL-13, which promotes transcriptional pathways leading to goblet cell expansion, shifts in the production of gel-forming mucins from MUC5B to the more pathologic MUC5AC (32), and alterations in antimicrobial peptides and ion transporters (4), resulting in the airflow obstruction and airway mucus plugging that are prominent features of type 2-high asthma (33). In the context of these dramatic changes in mucus quantity and quality, the expansion and programming of tuft cells that enables production of PGE2 may serve to hydrate the mucus and promote clearance via CFTR. The specific importance of CFTR activity in maintaining fluid transport in the allergic airway is reinforced by the observation that CFTR gene mutations are found in patients with asthma or nasal polyps (without cystic fibrosis) at higher rates than healthy control populations (34). While PGE2-enhanced mucus transport may be beneficial in the context of pathological mucus production in the allergic airway, further studies are needed to define the full spectrum of PGE2 effects, as well as the function of other secreted factors derived from tuft cells. Nevertheless, our study firmly establishes the IL-13-programmed tuft cell as part of the epithelial remodeling that occurs in the allergic airway.

While tuft cells display high expression of *PTGS1* (one of two enzymes that catalyzes the rate-limiting step in prostaglandin production), other cell types also produce PGE2. Some, such as mast cells, are concurrently elevated in nasal polyposis, allergic rhinitis, and asthma (35–37). Our data shows reduced PGE2 metabolites in epithelial organoids as well as whole tracheal tissue from tuft cell-deficient mice, suggesting that tuft cells are themselves a source of this molecule in airway epithelium. It is not clear how tuft cell-dependent PGE2 production differs qualitatively or quantitatively from that of other PGE2-producing cells. One possibility is that tuft cell-derived PGE2 acts predominantly on neighboring epithelial cells, while immune cell-derived PGE2 production may be more critical in the submucosa. Further, while our epithelial organoids strongly suggest tuft cells are themselves a source of PGE2, it is also possible that tuft cell-derived signals stimulate PGE2 production from other sources.

While PGE2 is known to be dysregulated in the type 2 inflammatory environment, its effects on cells and tissues have proven to be complex and pleomorphic. For instance, it can promote vasodilation and tissue edema through direct smooth muscle relaxation (38), has anti-inflammatory effects on both the innate and adaptive immune system (39), can promote stem cell survival and tissue regeneration (40), and has well-described function in gastrointestinal mucosal protection, secretion and motility (41). These functional consequences are context-dependent. While our work focuses on the epithelium, tuft cell-dependent PGE2 production likely has broad effects on diverse cell types in the respiratory tract.

Our finding that PGE2 activates CFTR-dependent epithelial ion channels in the respiratory epithelium to induce fluid secretion is consistent with a large body of literature showing similar action in the intestinal epithelium (27). While our study has focused on the role of PGE2 in epithelial ion transport and mucociliary function, it is notable that both the sinus and intestinal epithelium are susceptible to polypoid epithelial growth. In the intestine, polyp formation is dependent on local PGE2 production and can be blocked experimentally and clinically by inhibition of cyclooxygenase (COX)-1 or -2 (42, 43). Blockade of EP4 also abrogates intestinal polyp formation (44). Further, a number of the genes we find to be PGE2-responsive are linked to cell growth and differentiation and are dysregulated in some cancers (45–47), while a growing body of work has implicated tuft cells in cancer formation or progression (48–51). Despite these similarities, intestinal polyps are pre-cancerous while nasal polyps are stably benign. Nevertheless, in the context of this broad literature, one may hypothesize that tuft cell-derived products such as PGE2 could influence neoplastic growth including nasal polyp formation in a genetically susceptible host. Further study of the potentially pleiotropic impacts of tuft cell-dependent PGE2 production in the airway is needed.

The treatment paradigm for nasal polyposis, previously managed with surgery and topical corticosteroids, has been revolutionized by inhibition of IL-4 and IL-13. Yet the therapeutic effect of type 2 cytokine blockade is incomplete (52), suggesting additional pathologic mechanisms. Our data reveals an elevated PGE2 score in CRSwNP, raising the question of whether targeting this pathway may be of clinical benefit. While the patient group in this study was too small to correlate transcriptional signatures of type 2 inflammation, tuft cell activation, and PGE2 stimulation with clinical measures or disease endotypes, such correlations in larger patient cohorts will inform the use of these biomarker signatures for therapeutic discovery.

COX inhibitors have enjoyed success in the clinical management of intestinal polyposis, but such drugs prove problematic when deployed in nasal polyposis where a significant subpopulation display aspirin sensitivity (or aspirin exacerbated respiratory disease, AERD), characterized by wheezing and hives. The mechanism of this sensitivity to COX inhibition is thought to be due to shunting of arachidonic acid toward leukotrienes (LT) and reduced smooth muscle EP2 receptor activity leading to bronchoconstriction and inflammation (53). Reduced EP2 expression in AERD compared to aspirin-insensitive CRSwNP may further contribute to the relative imbalance of LT versus PGE2 effects on inflammation and bronchoconstriction (54). While the relative deficiency of PGE2 and excess LT mediated by COX inhibition is pathologic in AERD patients, pharmacologic manipulation of specific prostaglandin receptor subtypes could prove beneficial in CRS.

In sum, this work identifies a transcriptional shift in tuft cells in human airway disease that alters their signaling potential. It reveals a critical role for tuft cells in modulating epithelial homeostasis within the landscape of the respiratory tract and deepens our understanding of how epithelial functions are coordinately altered under allergic conditions.

## Methods

### Sinus Study Participants

Subjects between the ages of 18 and 75 years were recruited from the University of California, San Francisco Hospital (San Francisco, California) Otolaryngology clinic between 2013 and 2019 (Supplemental Table 1) to participate in a UCSF Sinus Tissue Bank. The UCSF Committee on Human Research approved the study, and all subjects provided written informed consent. Cytologic brushes were collected from nasal polyps or ethmoid sinus at the time of elective endoscopic sinus surgery from patients with physician-diagnosed chronic rhinosinusitis with or without nasal polyps on the basis of established guidelines. Subjects with cystic fibrosis were excluded from the study. Non-CRS control subjects were those who were undergoing endoscopic sinus surgery for alternative indications (i.e., septal deviation, pituitary surgery, etc.).

### Human Biospecimen Collection

Cytologic brushes obtained from the ethmoid sinus or nasal polyps were placed in RNAlater (Qiagen) for RNA extraction for bulk RNA sequencing, or into 10% FBS in RPMI (Gibco) on ice for single cell RNA-seq analysis or *in vitro* culture.

### Preparation of human biospecimens for single-cell RNA-seq

Brushes in 10% FBS/RPMI were vortexed on low speed for 2 minutes, brushes were removed and media was centrifuged at 300 g for 10 minutes at 4 degrees. Cell pellets were resuspended in 0.25% trypsin and placed on a thermomixer at 37°C at 300 rpm for 15 minutes. Trypsin was neutralized with complete media, and cells pelleted and then resuspended in 1x Pharmlyse (BD Bioscience) for 10 minutes followed by neutralization in complete media. Cells were pelleted at 300 g for 10 minutes and resuspended in 0.4% BSA in PBS prior to passing through a 40 μm cell strainer. 40,000 cells were loaded onto the 10X chip.

### Single-cell RNA-seq computational pipeline and analysis for human biospecimens

10X Single Cell 3’v3 chemistry was employed. Initial pre-processing of the 9 subjects’ 10X scRNA-seq data, including demultiplexing, alignment to the hg38 human genome, and UMI-based gene expression quantification, was performed using Cell Ranger (version 3.0, 10X Genomics).

### Preliminary Quality Control

Data were collected on 183,167 cells from 9 samples. We filtered out low quality cells with less than 200 genes detected or with greater than 75% of mapped reads originating from the mitochondrial genome. We additionally safeguarded against doublets by removing cells with a UMI count greater than the 98^th^ percentile of UMI counts for each sample. Prior to downstream analysis, select mitochondrial and ribosomal genes (genes beginning with MT-, MRPL, MRPS, RPL, or RPS) were removed. The preliminary quality-controlled dataset consisted of 176,803 cells and 23,778 genes.

To account for differences in coverage across cells, we normalized and variance stabilized UMI counts using the SCTransform method in the Seurat R package (55, 56). In addition to adjusting for sequencing depth, we also adjusted for the proportion of mitochondrial reads.

### Preliminary Analyses to Identify Epithelial Cells

#### Data Integration, Dimensionality Reduction, and Clustering

Data from the 9 subjects were combined using single cell integration implemented in Seurat v3. Integration was carried out using the top 30 dimensions from a canonical correlation analysis (CCA) based on SCTransform-normalized expression of the top 3,000 most informative genes, defined by gene dispersion using the Seurat’s SelectIntegrationFeatures function. Integrated data were then clustered and visualized using the top 20 principal components. For visualization, we reduced variation to two dimensions using Uniform Manifold Approximation and Projection (57) (UMAP; n.neighbors = 50, min.dist = 0.3). Unsupervised clustering was performed using a shared nearest neighbor (SNN) graph based on 20-nearest neighbors and then determining the number and composition of clusters using a smart local moving (SLM) algorithm (resolution = 0.4). This algorithm identified 20 preliminary clusters.

#### Cluster Markers

To identify cluster markers, we carried out pairwise differential expression analysis comparing log-normalized expression in each cluster to all others using a Wilcoxon rank sum test. Markers were identified as genes exhibiting significant upregulation when compared against all other clusters, defined by having a Bonferroni adjusted p-value < 0.05, a log fold change > 0.25, and >10% of cells with detectable expression. This analysis was then performed separately for each subject using Seurat’s FindConservedMarkers function to determine if marker genes were consistent across subjects. Cluster markers were required to have significant upregulation in at least half of the subjects.

Several clusters (11, 12, 13, 18, 19) had markers associated with both epithelial and immune cell types. These cells were sub-clustered using the same methods described above to separate immune and epithelial cells. Sub-clustering resulted in 8 epithelial and 7 immune sub-clusters. The dataset was divided into immune (8,153 cells) and epithelial (168,650 cells) datasets. Dimensionality reduction and clustering were performed separately for each dataset, resulting in 10 immune and 22 epithelial clusters.

#### Additional Quality Control and Doublet Detection

Potential doublets were assessed using the doubletCluster function in the scran R package (58, 59), which can be used to identify clusters of doublets, and the scds R package (60), which assigns a doublet score to each cell. Epithelial clusters 18, 20, and 21 were identified as likely doublet clusters and were removed from further analysis. In addition, cells with high binary classification-based doublet scores (BCDS >0.5) were also excluded from further analysis. Epithelial cluster 3 was comprised of cells with a high percentage of mitochondrial reads. Therefore, cluster 3 and all other cells with > 40% mitochondrial reads were excluded. Epithelial cluster 14 had only 3 consistent markers across subjects, including hemoglobin HBA1, HBA2 and HBB. These cells were also excluded from further analysis.

### Final Epithelial Data Set

The final quality controlled epithelial data set included 116,358 cells and 23,778 genes observed in at least 1 cell. Final data integration, dimensionality reduction, clustering and marker finding were performed, as described above. We identified 15 clusters, which were collapsed into 10 cell-types based on the expression of known marker genes (Supplemental Tables 2 and 3).

#### Sub-Clustering of Rare Epithelial Cells

Epithelial cluster 10 appeared to be a combination of rare cell types, including both ionocytes and tuft cells. We hierarchically clustered these cells based on scaled normalized expression of previously published cell type markers (19, 21, 22). Euclidian distance was used to measure similarity between cells, and cells were clustered using the complete linkage method in the hclust function in R.

Ionocyte and tuft cell markers were identified as described above. In addition, tuft cell markers were generated separately for control and polyp cells to understand heterogeneity in gene expression between polyp and control tuft cells. Tuft cell markers were categorized as “common” if they were significant markers in both control and polyp cells, “polyp specific” if they were significant in polyp cells only, and “control-specific” if they were significant in control cells only.

#### Pseudo-Bulk Differential Expression

To identify differentially expressed genes between control and polyp samples while accounting for clustering of cells within subjects, we performed pseudo-bulk differential expression analysis separately for each cell type (61–63). Within each cell type, expression counts were summed across all cells for a subject, resulting in a single expression measurement for each gene for each subject. Pseudo-bulk expression was compared between subjects with polyps and controls using bulk RNA-seq analysis methods with the edgeR R package (64, 65). Genes with Benjamini-Hochberg (66) adjusted p-values less than 0.05 were considered differentially expressed.

#### Hierarchical Clustering of Pan-Epithelial Genes

DEGs were classified as ‘pan-epithelial’ if they were up-regulated in polyp vs. control in 9 or more of the 11 cell types. These genes were hierarchically clustered based on Euclidean distance using the complete linkage method to identify modules of related genes.

#### Gene Signature Scores and Comparisons

Gene set signature scores were calculated for each cell by taking the average of scaled log-normalized expression of the genes in the signature set. We calculated the following gene signature scores: type 2 3-gene score (23), PGE2 score, common tuft marker score, polyp tuft marker score, Haber T1 and T2 scores (22), and Montoro T1 and T2 scores (6). Full lists of genes for each signature are available in Supplemental Table 5 or in listed references. Linear mixed models were used to compare gene signature scores between groups with the lmerTest R package (67, 68). Models included a subject-specific random intercept to account for clustering of cells within subjects. Tuft cell signatures from tuft or non-tuft cells from controls or polyps were also compared to published human tracheal tuft cells (19) or consensus mouse tuft cell markers(21).

### Comparison to Published Datasets

To confirm our scRNA-Seq findings, we re-analyzed data from Ordovas-Montanes et al (16). Quality control and filtering of cells was performed as described (16). Data integration, dimensionality reduction, clustering, marker finding and calculation and modeling of the PGE2 score were performed as described above. We identified 17 clusters, 7 of which included epithelial cells based on comparison to markers published in the original analysis. Gene scoring was performed as above. Statistical comparisons are included in Supplemental Table 7.

For analysis of bronchial epithelial brushes, we analyzed data from NCBI GEO under the accession number GSE109484. Normalized expression values were centered and scaled before calculation of gene signature (scores are described in “gene signatures scores and comparisons”). Signature scores were compared between groups using ANOVA.

### Gene Set Enrichment Analysis

Gene set enrichment analysis was performed using the EnrichR R package with the GO Biological Process 2018 database.

### RNA Extraction

Cytology brushes frozen in RNAlater were defrosted on ice and diluted with sterile 1x PBS. Samples were centrifuged at 18,000 g for 20 min and brushes were removed and placed into lyse E tube. Pellets were resuspended in RLT/BME and added to the Lyse E tube. Samples were agitated in a bead beater for 30 sec. Samples were centrifuged at 2000 rpm for 1 min and transferred to an Allprep (Qiagen) spin column. RNA and DNA were prepared according to the manufacturer’s instructions. Residual DNA was removed from the purified RNA by incubation with RNase-Free DNase (Promega) for 30 minutes at 37°C. DNase was removed from the preparation via a second RNA clean up using the Qiagen RNeasy Kit. RNA concentration was determined using Nanodrop (Thermo Scientific) and RNA quality was assessed using Agilent Pico RNA kit.

For cultured cells, ALI transwell inserts or organoids were lysed in RLT plus buffer (Qiagen) and RNA purified using the Qiagen RNeasy Kit according to the manufacturer’s instructions. Equal quantities of RNA were reverse transcribed using the SuperScript VILO cDNA synthesis kit (ThermoFisher) and amplified using Power SYBR Green PCR master mix (ThermoFisher) using the primers listed in Supplementary Table 4.

### Bulk RNA-seq for human biospecimens

For whole transcriptome sequencing we first used the Ion AmpliSeq™ Transcriptome Human Gene Expression Kit (Cat #A26325, Life Technologies) to enable gene-level expression analysis from small amounts of RNA. We generated barcoded sequencing libraries per the manufacturer’s protocol from 10 ng of RNA isolated from the 24 stimulation samples detailed above (12 pairs). Libraries were sequenced using the Ion PI template OT2 200 kit v3 for templating and the Ion PI sequencing 200 kit v3 kit for sequencing. Barcoding allowed all 24 samples to be loaded onto 3 PI sequencing chips and sequenced with an Ion Proton Sequencer using standard protocols. Read mapping was performed with the TMAP algorithm on the Proton server and read count tables for each gene amplicon generated using the Proton Ampliseq plugin. Read counts for gene amplicons across all 3 runs were merged to generate the final raw expression data.

### Bulk Gene Signature Scores and Comparisons

Expression data were normalized using the variance stabilizing transformation (VST) in the DESeq2 R package (69). Gene set signature scores were calculated for each sample by taking the average of scaled VST-normalized expression of the genes in the signature set (Supplemental Table 5). Linear regression models were used to compare gene signature scores between groups.

### Human respiratory epithelial cell culture

Human tracheal epithelial cells were harvested from deceased organ donors according to established protocols (70). Human sinus epithelial cells were harvested from research participants undergoing endoscopic surgery using a cytobrush as described above. For Ussing chamber measurements, the inferior turbinate of healthy individuals was sampled using a protocol approved by the National Jewish Health Institutional Review Board (HS-2832) and all donors provided written informed consent prior to the procedure. Tracheal, sinus, or nasal epithelial cells were seeded onto mitomycin-treated MRC5 or irradiated NIH/3T3 fibroblast feeder layers and cultured in cultured in conditional reprogramming culture (CRC) medium (71) supplemented with the ROCK inhibitor Y-27632 (ApexBIO). Expanded cells were plated on collagen-coated Transwell inserts (Corning) cultured with Pneumacult ALI (StemCell) for 21-28 days according to the manufacturer’s instructions.

For organoid culture, expanded basal cells were plated in matrix (80% Matrigel, 20% media) in Pneumacult Airway Organoid Media (StemCell) according to manufacturer’s instructions for 21-24 days. Cells were stimulated with PGE2 (Sigma, 1 µg/ml) daily, EP2 inhibitor (EP2_inh_, PF-04418948, Cayman Chemicals, 10 µM), EP4 inhibitor (EP4_inh_, L-161,982, Cayman Chemicals 10 μM), CFTR inhibitor 172 (CFTR_inh_, Sigma, 10 µM).

### Ussing chamber

Electrophysiological analyses were performed in an Ussing Chamber (Physiologic Instruments, San Diego, CA). Epithelia on Transwell inserts were mounted in an Ussing chamber and bathed in a modified Ringer’s solution (120 mM NaCl, 10 mM D-Glucose, 3.3 mM KH_2_PO_4_, 0.83 mM K_2_HPO_4_, 1.2 mM MgCl_2_, 1.2 mM CaCl_2_, 25 mM NaHCO_3_, pH 7.4), maintained at 37°C and gassed with 5% CO_2_/95% O_2_. Epithelia were analyzed under short-circuit conditions with intermittent pulsing (200 ms pulses at +/- 5 mV). Cultures were treated acutely in the Ussing chamber with subsequent additions of apical amiloride (100 µM, Alfa Aesar), apical and basolateral forskolin (20 µM, Tocris) and IBMX (100 µM, Sigma) (F/I), apical CFTR(inh)-172 (10 µM, CFTR Chemical Compound Distribution Program) and apical ATP (100 µM, Sigma). Inhibition of epithelial sodium channel-dependent current with amiloride enabled normalization of basal currents between epithelial donors.

### Mice

All experimental procedures on mice were approved by the UCSF Animal Care and Use Committee. IL-25 reporter (IL25^F25^) (10) and *Pou2f3*^-/-^ (72) mice on a B6 background have been described. For single cell sequencing, male C57BL/6J mice were obtained from Jackson Laboratories (stock # 000664) at 9 weeks of age and maintained under specific pathogen free conditions with 12 hr light/day cycle, and ad libitum access to food and water. Following 1 week of acclimation to our facility, mice were rapidly injected by tail vein injection with 2 µg of pLive *in vivo* expression vector (Mirus Bio) into which the mouse IL-13 coding sequence was cloned, in a volume of sterile PBS equivalent to 10% of body weight, as described (72). Control mice received an injection of IgG1 expression vector. Overexpression of IL-13 was verified by measurement of serum IL-13 using a mouse IL-13 enhanced sensitivity flex set (BD Biosciences), analyzed on an LSR Fortessa flow cytometer (BD) with FCAP Array software (BD) 1 week later, or at the time of sacrifice. For PGE2 measurements or organoid culture, the protocol was as described, except that *Pou2f3^-/-^* or WT mice were bred at UCSF, and age and sex-matched males and females between 6-10 weeks of age were used.

### Mouse nasal epithelial preparation for single cell sequencing

4 weeks after initial plasmid injection, 3 IL-13-overexpressing mice and 4 controls were sacrificed and the nasal epithelium dissected. Tissue was minced with scissors and epithelium separated to single cell suspension by incubation in HBSS containing 5 mg/mL Dispase II (Gibco) for 45 minutes at room temperature, followed by 15 minutes in HBSS containing 25 µg/mL DNAse I (Roche), and filtration through a 70 μm strainer. Following lysis of red blood cells, cells were stained for CD45 (clone 30-F11, BioLegend Cat #103108) and EpCAM (clone G8.8, BioLegend Cat #118233) antibodies for 30 minutes on ice, followed by staining with 4′,6-diamidino-2-phenylindole (DAPI) to identify dead cells. Live, CD45-cells were sorted into RPMI media on a MoFlo sorter, pooled in equal proportion from like mouse samples, and then resuspended in 0.4% BSA in PBS before submission of 30,000 cells per sample for sequencing. A small aliquot of cell suspension was reserved for analysis on an LSR Fortessa flow cytometer, and demonstrated >90% viability (by DAPI exclusion) and absence of CD45+ cells.

### Single-cell RNA-seq computational pipeline and analysis for mice

Single cell libraries from 30,000 cells per sample were prepared with the Chromium Single Cell 3ʹ GEM, Library & Gel Bead Kit v3 (10 x Genomics PN-1000075) following the manufacturer’s protocol. The libraries were sequenced on the NovaSeq 6000 at the UCSF Institute for Human Genetics. Initial pre-processing including demultiplexing, alignment to the mouse genome, and UMI-based gene expression quantification, was performed using Cell Ranger (version 3.0, 10X Genomics).

Data from pooled samples from 3-4 mice per treatment group FACS sorted to deplete immune cells (as described above) were integrated before filtering out low quality cells according to the following parameters: min.cell =3, min.features=200; nFeature_RNA >= 300; nFeature_RNA< 5000; nCount_RNA < 20000; percent mitochondrial reads < 15. 14,172 filtered cells were used for analysis. We normalized and variance stabilized UMI counts using the SCTransform method in the Seurat R package (55, 56), adjusting for the proportion of mitochondrial reads. Cluster markers were manually compared to published consensus tuft cell gene expression (21) to identify a single cluster of tuft cells. We subset only this cluster and re-ran dimensionality reduction and clustering using SCTransform, then removed contaminating non-tuft cells before the final subclustering. Diffuse expression of canonical markers including *Pou2f3*, *Avil*, *Trpm5* and *IL17rb* and absent expression of other non-tuft lineage defining markers confirmed the purity of the final Seurat object. We merged resulting clusters of tuft cells and then further subset and re-ran dimensionality reduction and clustering on the tuft cells alone. Marker lists are provided in Supplemental Table 6. Linear regression models were used to compare the polyp tuft score between tuft cell subsets..

### Mouse tracheal epithelial organoid culture

Organoid growth media was comprised of DMEM-F12 (Gibco) supplemented with 10mM HEPES (Gibco), 1x Glutamax (Gibco), 100 U/mL Penicillin-Streptomycin, 1mM N-acetyl cysteine (Sigma), 1x N2 and B27 supplements (Gibco), 0.5 μg/mL mouse R-spondin 1(Peprotech), 100ng/mL mouse Noggin (Peprotech), 20 ng/mL mouse epidermal growth factor, 25 ng/mL FGF2 (Peprotech), 100 ng/mL mouse FGF10 (Peprotech) and 10 μM Y-27632 (Cayman Chemical).

Mouse tracheas of indicated genotypes were removed, cleaned, and fileted before digestion at 37°C in HBSS containing 5mg/mL dispase II (Sigma). After vigorous vortexing, the cells and partially-digested tracheal tissue were pelleted by centrifugation and then incubated for an additional 15 minutes with HBSS containing 25 µg/mL DNAse I (Roche). After additional vortexing, cells and remaining tissue were filtered through a 40 µm mesh, pelleted, and contaminating red blood cells lysed using Pharmlyse buffer (BD). Cells were resuspended in organoid growth media with the addition of 10% fetal bovine serum, 2.5 μg/mL amphotericin B (Gibco), 100 U/mL nystatin and 50 μg/mL gentamicin on 10 mm dishes coated with rat tail collagen (Sigma). Confluent cells lifted with 0.25% trypsin for 30 minutes before seeding into 50 µl of matrix (80% Matrigel, 20% media). Media was collected on day 8 of culture, centrifuged at 300g for 10 minutes to remove cellular material and debris, and stored at −80°C. Aliquots were analyzed on Prostaglandin E2 Parameter Assay Kit (R&D Systems) according to manufacturer’s instructions.

### PGE2 metabolite (PGEM) measurement

Mouse tracheas were cleaned, rinsed in cold PBS containing 7.5 µg/mL indomethacin and snap frozen. Frozen samples were homogenized in indomethacin-containing PBS and debris removed by centrifugation. Lysates underwent acetone precipitation, derivation and assayed according to manufacturer’s instructions for Prostaglandin E Metabolite ELISA Kit (Cayman Chemicals).

### Imaging

Organoids were imaged on an inverted Nikon A1R with DS-Fi3 camera or upright light microscope (Leica). Manual measurements of organoids at estimated maximal diameter were performed on NIS Elements or Fiji software.

For immunofluorescence of mouse tissue, 4% PFA-fixed, sucrose-protected anterior skulls were embedded in optimal cutting media (OCT, Sakura) and cryosectioned at 8 µm thickness midway between the incisors and nares. For whole mount tracheas, entire fixed tracheas were fileted open and cleaned of attached connective tissue. Tissues were stained for DCLK1 (Abcam Cat# ab31704) and RFP (SicGen, Cat# AB8181) followed by appropriate secondaries, counterstained with DAPI, mounted, and analyzed on a Nikon A1R confocal microscope using NIS Elements (Nikon) and Fiji (ImageJ) software.

For particle transport measurements, 20 μL of 5 μm fluorescently-labelled polystyrene beads (Bang’s Laboratories) in PBS were applied to the center of the apical surface of human tracheal epithelia on Transwell inserts and PGE2 (1 µg/ml) +/- CFTR inhibitor 172 (10 µM) applied on the basolateral side. After 1 hour in a standard tissue culture incubator, tissue culture plates were transferred to a 37°C, 5% CO_2_-injected incubated stage of a Nikon A1R confocal microscope and 30 second videos obtained for each well. Track speeds for individual particles were calculated using Imaris 9.7.2 software. Track speeds were log transformed to obtain a normal data distribution before testing by ANOVA.

### RNAscope Multiplex Fluorescence in situ hybridization

Human sinus tissues were fixed in 10% neutral buffered formalin and embedded in paraffin. Paraffin blocks were sectioned onto glass microscope slides, dried overnight, and baked for 1 h at 60 °C before immediately proceeding with the RNAScope Multiplex Fluorescent v2 assay according to the manufacturer’s protocol (Advanced Cell Diagnostics, ACD). Deparaffinization was performed with 100% Xylene followed by 100% Ethanol. The tissue sections were pretreated with RNAscope Hydrogen Peroxide, followed by target retrieval and Protease plus pretreatment. Pretreated sections were hybridized with target probes *POU2F3* or *FOXI1* for 2h at 40 °C. Hybridized slides were left overnight in 5XSSC before proceeding with the amplification and labeling steps. Targeted probes were labeled with Opal dyes (1:500, Akaya Biosciences) and cell-cell junction were visualized with E-Cadherin antibody (1:2,000; ECM biosciences Cat # CM1681, Clone # M168) followed by Alexa fluorochrome-conjugated donkey anti-Mouse IgG (1:500, ThermoFisher, Cat # A32787). For nuclear staining, sections were incubated with DAPI for 5min at room temperature. All stained sections were mounted with ProLong Diamond Mount Medium (Invitrogen) and imaged on an Echo Revolve R4 fluorescence microscope.

### Data and Materials availability

Single cell RNA-seq data have been deposited at the National Center for Biotechnology Information/Gene Expression Omnibus (GEO) under accession number GSE202100. Code used to carry out data analysis is available upon request.

### Statistics

Statistical methods used for single cell and bulk sequencing analysis of human subjects is described in detail in the corresponding sections of the methods and figure legends. Specific statistical comparisons (with corrections for multiple comparisons) relating to Figure 2 are included in Supplemental Table 7. For mouse studies, a power calculation was made whenever possible based on pilot studies to determine the number of mice needed to detect what we considered to be a biologically meaningful difference. For mouse studies, t-testing or one-way ANOVA was performed in GraphPad Prism, or non-parametric tests as indicated in figure legends.

### Study Approval

Sampling from human subjects was approved by the UCSF Committee on Human Research, and all subjects provided written informed consent. All experimental procedures on mice were approved by the UCSF Animal Care and Use Committee.

## Author contributions

All authors substantially contributed to the acquisition, analysis or interpretation of data for the manuscript and drafting, revising and critically reviewing the manuscript for important intellectual content. MEK, EKC, RML, MAS, and EDG conceptualized the experiments. JGG, SDP, ANG recruited subjects and obtained epithelial brushes. MEK, CMM, RA, SY, MTM, CAS, IZ, YL, KL and EDG performed experiments and analyzed data. MEK, CMM, MAS, and EDG synthesized the data and drafted the manuscript, which was further edited by MEK, CMM, RML, MAS, and EDG. All authors approved the final version of this manuscript.

## Acknowledgements

The authors would like to acknowledge Dr. Robert Bridges, the Rosalind Franklin University of Medicine, and the Cystic Fibrosis Foundation for the contribution of compounds to this work through the CFTR Chemical Compound Distribution Program. We thank Kyle Marchuk of the Biological Imaging Development CoLab at UCSF Parnassus for training and support for video microscopy and analysis using Imaris. Technical support for single cell sequencing was provided by the UCSF Institute for Human Genomics. We are grateful to the financial support of the National Institutes of Health (5P01HL107202-07 to RML and MAS; R01AI026918 to RML; 5R01AI136962 to EDG; R01HL128439, R01HL135156, and P01HL132821 to MAS; F32HL140868 and T32HL007185 to MEK; F32HL158174 to CAS), the Department of Defense (MAS), the Howard Hughes Medical Institute (RML), the A.P. Giannini Foundation (MEK), the Webb-Waring Early Career Investigator Award from the Boettcher Foundation (CMM), the Nina Ireland Program for Lung Health at UCSF (EDG), the UCSF John A. Watson Faculty Scholar Fund (JGG), the SABRE Center at UCSF (RML, EDG), the Cystic Fibrosis Foundation (PEB and PLZ), and the Eugene F. and Easton M. Crawford Charitable Lead Unitrust (CAS and PEB). This work would not have been possible without the support and participation of our patients.

