## Supplementary Data for "IL-13-programmed airway tuft cells produce PGE2, which promotes CFTR-dependent mucociliary function"

**Supplemental Data**

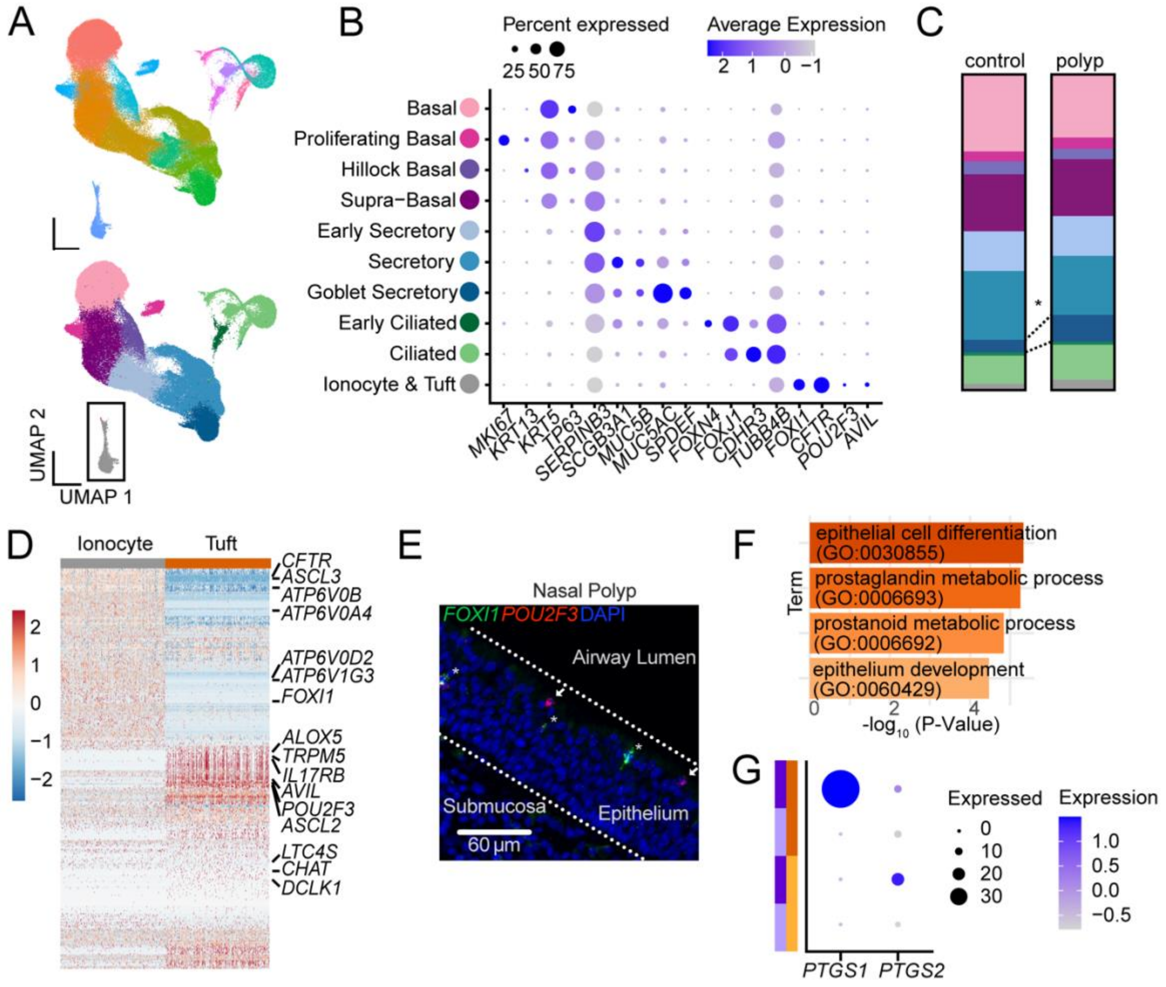


**Supplemental Figure 1. Single cell sequencing from human nasal polyps reveals a novel tuft cell phenotype that is unique to human nasal polyps.**

**(A)** Uniform manifold approximation and projection (UMAP) of scRNA-seq of epithelial cells from healthy ethmoid sinus (control; n=4) or nasal polyp (n=5) (total cells = 116,358) colored according to 15 clusters identified by Seurat (top panel) or after aggregation into 10 cell types (bottom panel). **(B)** Representative marker genes for each of the cell types. **(C)** Percent of each cell type among total epithelial cells. *p<0.05 by Mann-Whitney t test with Sidak correction for multiple comparisons. all other cell type comparisons non-significant. **(D)** Hierarchical sub-clustering of tuft cells and ionocytes. **(E)** RNA *in situ* hybridization for *FOXI1* (green) and *POU2F3* (red) identifies ionocytes (asterisk) and tuft cells (arrow) within nasal polyp epithelium. **(F)** Enriched pathways among differentially expressed genes between tuft cells from polyps versus controls. (**G**) Polyp tuft cells highly and uniquely express *PTGS1*.

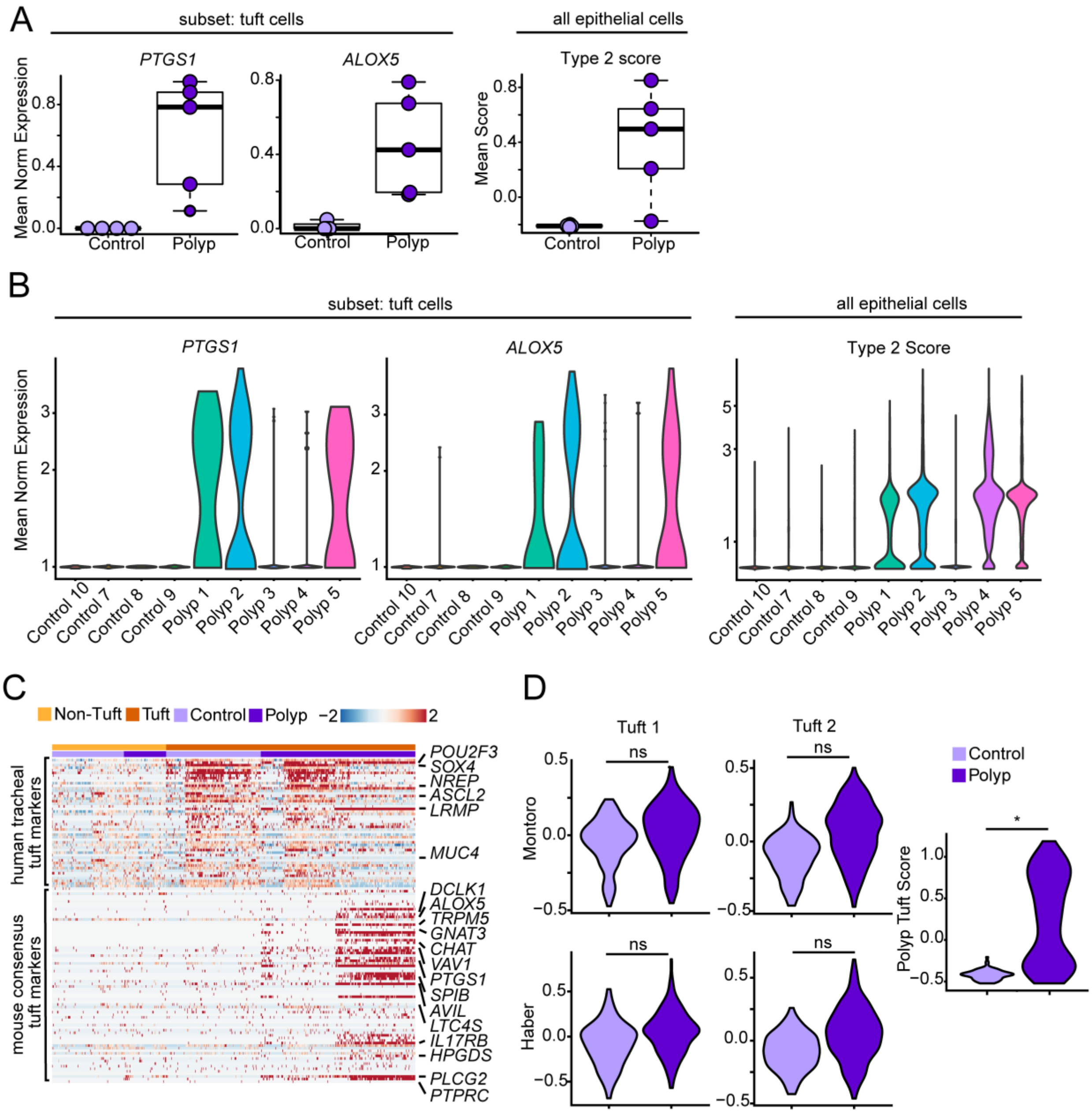


**Supplemental Figure 2. Heterogeneity of allergic tuft cell activation is distinct from previously-described tuft cell heterogeneity.**

**(A)** Mean normalized expression of *PTGS1* and *ALOX5* in tuft cell subset and Type 2 score in total epithelium between subjects. (**B**) Mean normalized expression of *PTGS1* and *ALOX5* in tuft cell subset and Type 2 score in total epithelium by subject. (**C**) Published markers of human tracheal tuft cells and consensus mouse tuft cell markers in tuft or non-tuft cells from control or polyp sinus brushes. **(D)** Scoring of published markers of “Tuft 1” or “Tuft 2” subsets from published work, or our polyp tuft score in control or polyp tuft cells. *$p<0.05$ by linear mixed model.

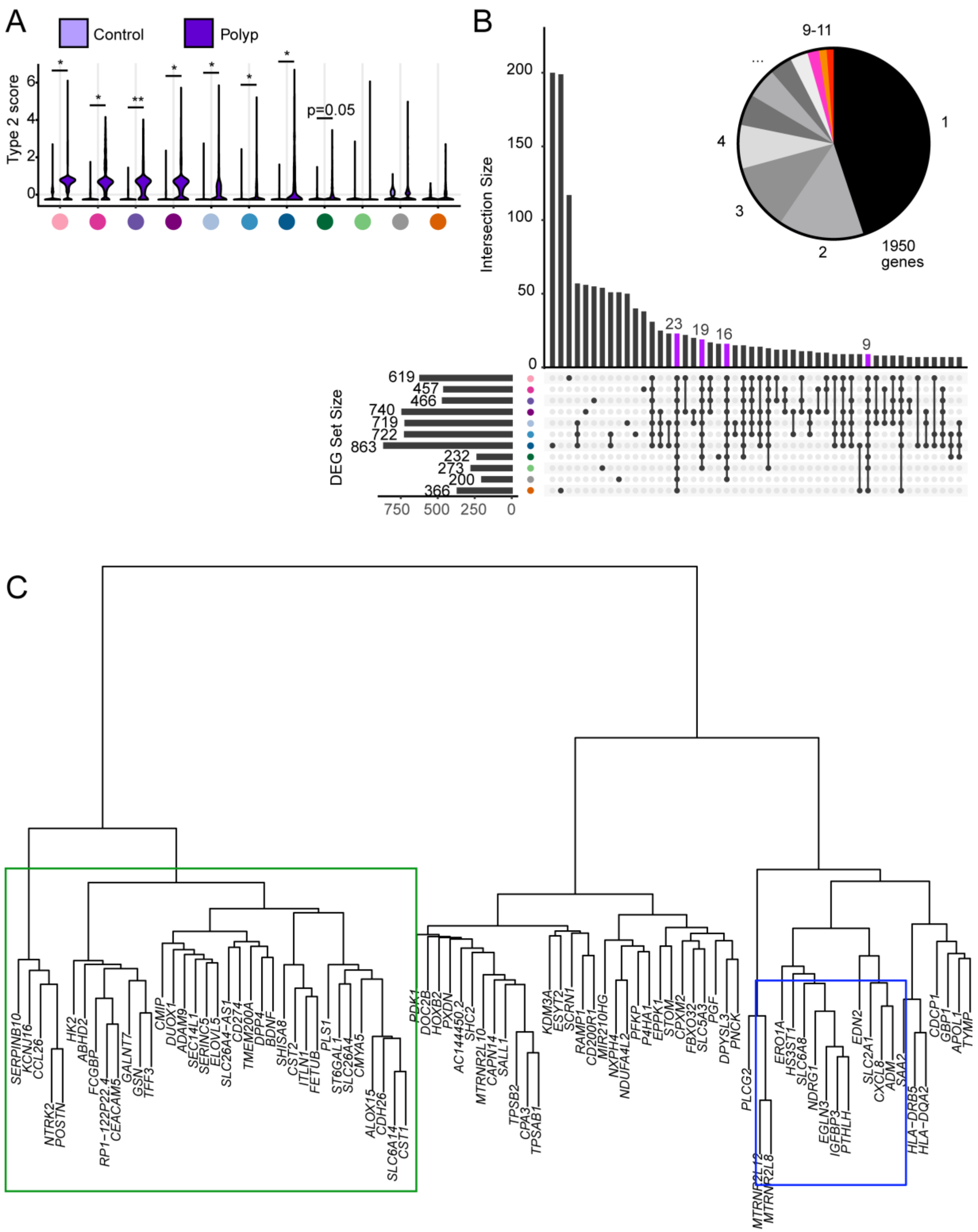


**Supplemental Figure 3. Pan-epithelial transcriptional analysis of nasal polyps.**

**(A)** Type 2 inflammatory 3 gene score in each cell type in control (light purple) or polyp epithelium (dark purple). *p<0.05; **p<0.01; ***p<0.001; ****p<0.0001 by linear mixed model. (**B**) Upset plot showing DEGs from each cell type and overlap (intersection) of DEGs between cell types. Pan-epithelial DEGs highlighted in violet. Inset pie chart indicates the proportion of total identified DEGs that were upregulated in multiple cell types; DEGs that were commonly upregulated in 9, 10, or 11 cell types are highlighted in pink, orange, and red, respectively, while DEGs upregulated across fewer distinct cell types are colored in shades of grey. **(C)** Hierarchical clustering of pan-epithelial up-DEGs (polyp > control), highlighting modules of genes with similar behavior. Genes recognized *a priori* and confirmed to be IL-13-responsive are outlined in green, while the PGE2 response module is outlined in blue.

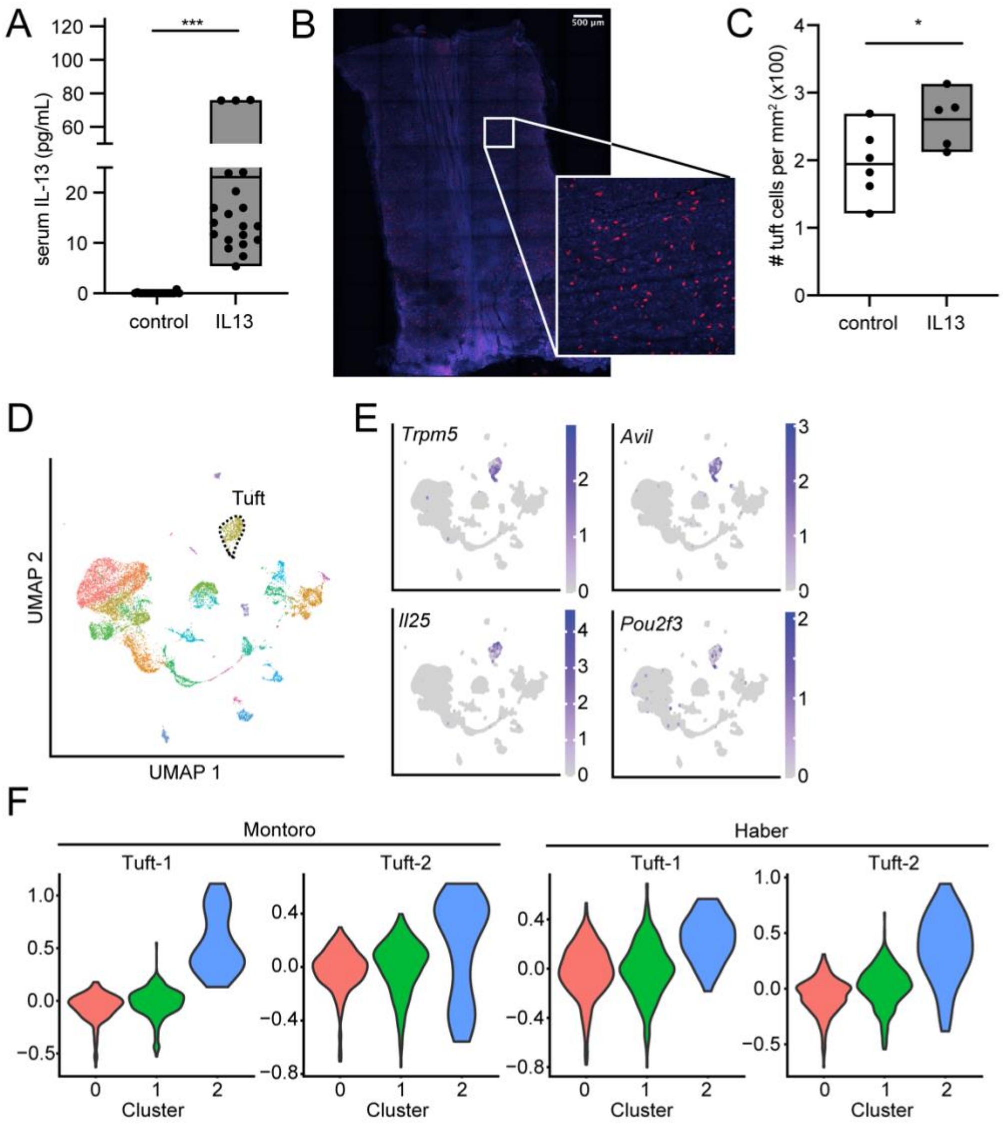


**Supplemental Figure 4. Overexpression of IL-13 increases airway tuft cells in mice.**

**(A)** Serum IL-13 concentration in IL-13-plasmid-injected or control plasmid injected mice. n=20 mice/group pooled from 2 experiments, and representative of measurements from $\geq$ 10 individual experiments. ***P<0.001 by t-test **(B)** Representative whole mount of mouse trachea used to enumerate tuft cells. Tuft cells marked by staining of DCLK1 (red), nuclei marked by DAPI (blue). Bar and inset represent 500 µm. **(C)** Quantification of tuft cells from whole mounted traches of IL-13-treated or control mice. n=5-6 mice per group, p<0.05 by t-test. **(D)**

UMAP of single cell sequencing of nasal epithelium from mice, with tuft cells outlined with dotted line. **(E)** Expression of tuft cell markers. The highlighted and outlined cluster was used to further analyze mouse tuft cells as in Figure 3. (**F**) Scoring of markers of “Tuft 1” or “Tuft 2” subsets from published work across our mouse nasal tuft cell subsets.

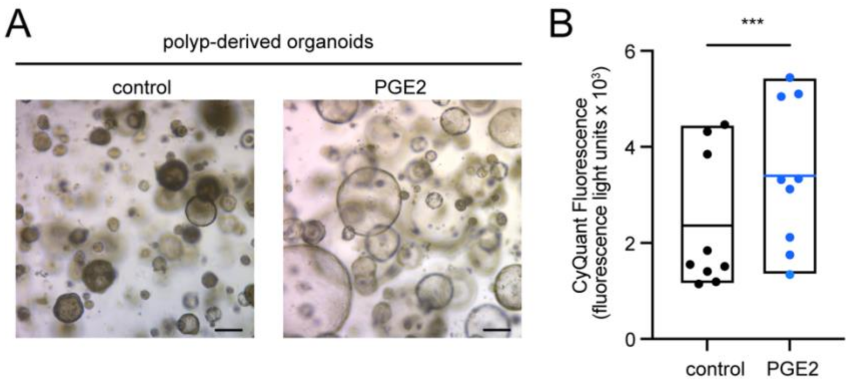


**Supplemental Figure 5. PGE2 induces swelling and increases cellularity in airway epithelial organoid culture.**

(**A**) Representative image of organoids grown from polyp epithelium. Scale bars indicate 250 µm. (**B**) Cell number as reflected in total DNA content of organoids as in Figure 5. pooled from 3 similar experiments from different epithelial donors, each treatment type in triplicate wells. Bars, mean +/- min/max. ***p<0.001 by t-test.

**Supplemental Table 1. Characteristics of subjects sampled for sequencing.**

1a. Patients with Epithelial Brushes for Single Cell RNA Sequencing

| | Control 7 | Control 8 | Control 9 | Control 10 | Polyp 1 | Polyp 2 | Polyp 3 | Polyp 4 | Polyp 5 |
|---|---|---|---|---|---|---|---|---|---|
| Age | 34 | 34 | 25 | 54 | 47 | 37 | 65 | 45 | 68 |
| Sex | M | F | F | M | F | F | F | F | F |
| Race | Hispanic | White NH | White NH | White NH | Hispanic | White NH | White NH | White NH | White NH |
| Smoking | No | No | No | Current | No | No | Former | No | No |
| Topical Steroids | No | No | No | No | No | Yes | Yes | No | No |
| Oral Steroids | No | No | No | No | No | No | No | No | No |
| Antibiotics | No | No | No | No | No | No | No | No | No |

1b. Patients with Epithelial Brushes for Bulk RNA Sequencing

| **Characteristic** | **Control (n=8)** | **CRSsNP (n=7)** | **Polyp (n=24)** | **p value Control vs Polyp** |
|---|---|---|---|---|
| Age (y) | 28.0 (12.3) | 54.9 (24.7)** | 49.1 (13.7)** | p=0.006 |
| Female sex, no. (%) | 1 (12.5) | 3 (42.9) | 10 (41.7) | |
| Race, no (%) | | | | |
| White Non-Hispanic | 3 (37.5) | 6 (85.7) | 18 (75.0) | |
| White Hispanic | 1 (12.5) | 0 (0) | 1 (4.2) | |
| Asian | 1 (12.5) | 1 (14.3) | 2 (8.3) | |
| African American | 0 (0) | 0 (0) | 2 (8.3) | |
| Mixed/Other | 3 (37.5) | 0 (0) | 1 (4.2) | |
| Current or Former Smoker, no (%) | 1 (12.5) | 4 (57.1) | 7 (29.2) | |
| Receiving ICS, no (%) | 0 (0) | 0 (0) | 21 (87.5)**** | P<0.0001 |
| Receiving oral steroids, no (%) | 0 (0) | 6 (85.7)*** | 20 (83.3)**** | P<0.0001 |
| Receiving Antibiotics, no (%) | 0 (0) | 7 (100)*** | 7 (29.2) | |

* indicates statistically significantly different from controls

**Supplemental Table 2. Percentage of cells of each cluster or type in each subject sample.**

(multi-tab spreadsheet in auxiliary file)

**Supplemental Table 3. Marker lists used to identify cell types.**

(multi-tab spreadsheet included as auxiliary file)

**Supplemental Table 4. Quantitative PCR primers.**

| Gene Symbol | Forward | Reverse |
|---|---|---|
| *PTHLH* | GAAACGCAAGGAGCAGGAAA | CCCAGTCACTCCAGAGTCTAAC |
| *RPL13A* | AACCCTGCGACAAAACCTCCTC | CAGAAATGTTGATGCCTTCACAGC |
| *SLC6A8* | GGCCTGGGGCTTCTATTACC | CAGCCTCAAGACTTTGTTCTCC |
| *CST1* | GGGTGAATTACTTCTTCGACGTAGA | CACAACTGTTTCTTCTGCAGTTCTG |
| *POSTN* | CTCATAGTCGTATCAGGGGTCG | ACACAGTCGTTTTCTGTCCAC |
| *NTRK2* | CCTGTAAAGCGGTTCGCTATG | ACCTTATCCAGGACGACATCCCTA |
| *ALOX15* | CCAACCACCAAGGATGCAA | TCTGCCCAGCTGCCAAGT |
| *IGFBP3* | AGCACAGCACCCAGACTTCA | GGTCGGCCGCTTCGA |
| *NDRG1* | CCAACAAAGACCACTCTCCTC | CCATGCCCTGCACGAAGTA |
| *EGLN3* | CTGGGCAAATACTACGTCAAGG | GACCATCACCGTTGGGGTT |
| *ERO1A* | CCTGAGCGCTACACTGGTTAC | TCTCTTCACTTGTCCCTTGACC |

**Supplemental Table 5. Gene lists used for scoring.**

(multi-tab spreadsheet included as auxiliary file)

**Supplemental Table 6. Mouse tuft cell marker list.**

(multi-tab spreadsheet included as auxiliary file)

**Supplemental Table 7. Statistical analyses relating to Figure 2 with corrections for multiple comparisons.**

(multi-tab spreadsheet included as auxiliary file)
